# Single-cell analysis of patient-derived PDAC organoids reveals cell state heterogeneity and a conserved developmental hierarchy

**DOI:** 10.1101/2020.08.23.263160

**Authors:** Teresa G Krieger, Solange Le Blanc, Julia Jabs, Foo Wei Ten, Naveed Ishaque, Katharina Jechow, Olivia Debnath, Carl-Stephan Leonhardt, Anamika Giri, Roland Eils, Oliver Strobel, Christian Conrad

## Abstract

Pancreatic ductal adenocarcinoma (PDAC) is projected to be the second leading cause of cancer mortality by 2030. Bulk transcriptomic analyses have distinguished ‘classical’ pancreatic tumors from ‘basal-like’ tumors with more aggressive clinical behaviour. We derived PDAC organoids from primary tumors of 18 patients, together with two matched samples from liver metastases. By single-cell RNA sequencing, we show that PDAC organoids consist of ductal cells with patient-specific expression of several gene groups, including genes which encode cell surface proteins. We report ‘classical’ and ‘basal-like’ cells coexisting within single primary tumors or metastases, with greater intratumor subtype heterogeneity linked to higher tumor grade. Single-cell transcriptome analysis of PDAC organoids and primary PDAC identified distinct tumor cell states shared across patients, including a cycling progenitor cell state and a differentiated secretory state. We show that these cell states are connected by a differentiation hierarchy, with ‘classical’ subtype cells concentrated at the endpoint of this hierarchy. In an imaging-based drug screen, expression of ‘classical’ subtype genes also correlates with better response to clinical drugs. Our results thus uncover a functional hierarchy of PDAC cell states linked to transcriptional tumor subtypes, and support the use of PDAC organoids as a clinically relevant model for *in vitro* studies of tumor heterogeneity.

## Introduction

Pancreatic ductal adenocarcinoma (PDAC) is the most common pancreatic cancer type ^1^, with a current 5-year survival rate of just 9% ^2^, and is predicted to be the second leading cause of cancer mortality by 2030 ^3^. Only 10–20% of PDAC are resectable at diagnosis, and response to chemotherapy is generally poor although long-term survival is achieved in a small fraction of patients ^4, 5^.

Large-scale genomic studies have identified recurrent genetic alterations in PDAC, with KRAS driver mutations detected in over 90% and inactivating mutations or deletions of TP53, SMAD4, and CDKN2A in over 50% of tumors ^6–9^. These alterations converge onto a limited number of aberrant signalling pathways ^10, 11^. Metastatic lesions have been shown to share identical driver gene mutations with the primary tumor ^12^, further supporting a high level of genomic uniformity in PDAC. Defining the origins of clinical heterogeneity in PDAC to enable patient stratification and appropriate patient-specific treatment choices therefore remains a key challenge.

Transcriptomic analyses of PDAC have resulted in several subtype classification schemes ^11, 13, 14^, with most evidence supporting a distinction between ‘classical’ and ‘basal-like’ tumors ^15^. The ‘classical’ subtype is associated with a higher level of mucinous features in histopathological assessment and longer survival ^14^. Whether these subtypes reflect genetically distinct cells, different evolutionary pathways, or different progression status, remains unclear.

Whereas most earlier studies of transcriptional features of human PDAC were limited to a single sample per patient, recent RNA sequencing data from multiregion-sampled biopsies or even single tumor cells demonstrate a previously unappreciated degree of intratumoral heterogeneity in PDAC. In particular, transcriptionally and histologically defined subpopulations exhibiting ‘classical’ or ‘basal-like’ features were found to coexist in metastases from the same patient or even within the same tumor sample ^16^. Furthermore, evidence from multiregion-sampled metastatic pancreatic cancers suggests that basal-like cell populations may emerge as a subclonal population within classical PDAC tumors ^17^.

Regardless of sampling resolution, attempts to delineate transcriptional tumor subtypes or distinct functional cell types in primary PDAC are often confounded by differences in neoplastic cellularity, resulting in the erroneous inclusion of transcriptional features present in stromal or normal pancreas cells ^18^. Differences in cell type composition were also described in recent single-cell RNA sequencing (scRNA-seq) efforts, which identified distinct populations of cells within surgically resected PDAC; these include including immune cells, fibroblasts and endothelial cells as well as abnormal and malignant ductal cells ^19, 20^. To simplify the study of PDAC tumor cells, patient-derived organoid models of human PDAC were introduced within the past five years, which reflect histopathologic, proteomic, genomic and transcriptomic features of the original tumors yet remain experimentally tractable ^21–23^. Such models offer great potential for detailed analyses of PDAC biology and developing therapeutic approaches.

To investigate the functional identity and hierarchical relationships of PDAC cells, we here performed scRNA-seq of PDAC organoids of primary tumors and metastatic samples from 18 patients. We show that ‘classical’ and ‘basal-like’ cells may coexist within the same sample, and the level of subtype heterogeneity is linked to tumor grade. Despite transcriptional differences between tumors, patient-derived organoids share functional tumor cell states that are connected by a differentiation hierarchy also present in primary PDAC samples. Our results support the use of PDAC organoids to model tumor heterogeneity *in vitro*.

## Results

### PDAC organoids comprise malignant ductal cells

To enable the *in vitro* study of human PDAC, we derived 24 tumor organoid lines from samples taken during surgery. 18 samples were obtained from primary tumors from individual patients. In one case, distinct organoid lines were derived from two pancreatic sites within the same primary tumor, providing a biological replicate (p080 and p081), and in another case, a technical replicate was generated by analysing one organoid line at different passage numbers (p039 and p039b). We also obtained samples of two different liver metastases in addition to the pancreatic primary tumor from one patient (p083, p084 and p085), and two samples from unmatched metastases (Supplementary Table 1 and Figure 1a,b). All tumors were classified as PDAC based on histological assessment. Patient-derived PDAC organoids, as well as nine of the biopsy samples, were classified based on bulk RNA sequencing as either basal-like or classical PDAC according to the subtypes defined by Moffitt *et al.* ^14^; in each case, the PDAC organoid subtype corresponded to the biopsy sample subtype (Methods and Supplementary Table 1).

**Figure 1:**
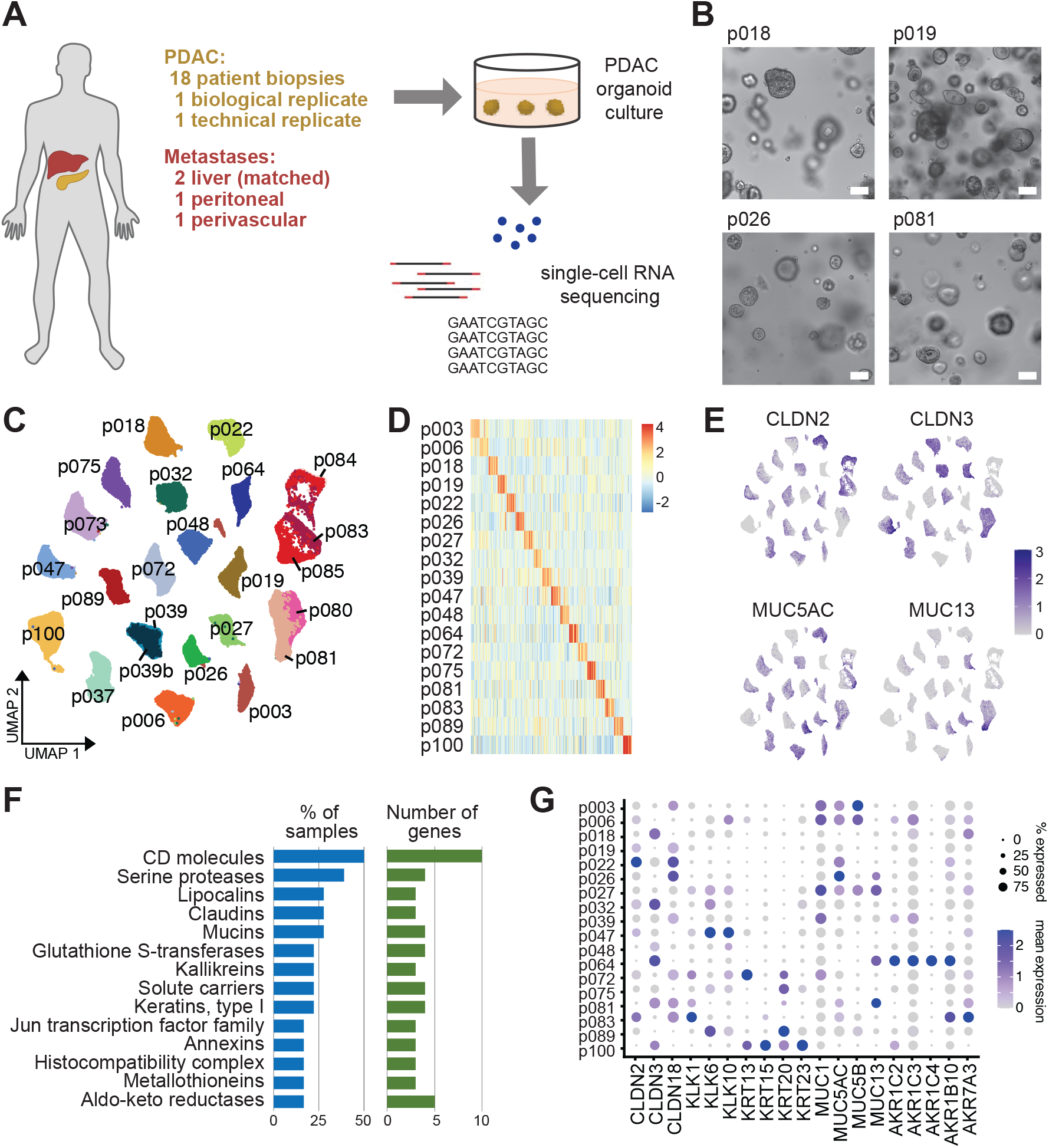
Intertumor heterogeneity in patient-derived PDAC organoids. **A)** Schematic of the experimental workflow. **B)** Example images of four patient-derived PDAC organoid lines after 10 days of culture. Scale bars, 100 µm. **C)** UMAP embedding of 24 scRNA-seq samples from 18 patients, showing that cells cluster by patient origin (p039 is a technical replicate of p039b, p080 is a biological replicate of p081, p084 and p085 derive from liver metastases matched to p083). **D)** Heatmap showing the expression of the 20 most highly differentially expressed genes per patient. **E)** Expression of selected patient-specific genes, using the same UMAP representation as in (C). **F)** Gene group analysis of the 20 most highly differentially expressed genes per patient. Bar plots show the proportion of patients in which the indicated gene groups were overrepresented (blue), and the number of unique genes from each gene group (green). **G)** Expression of selected patient-specific genes across all patients.

Single-cell RNA sequencing of all 24 tumor organoid lines resulted in transcriptomic information for a total of 93,096 cells after quality control, with a median of 3,877 cells from each organoid line and a median of 4,162 genes detected per cell (Methods and Supplementary Figure 1A).

To determine the cell type identity of PDAC organoid cells, we compared single-cell transcriptomes from PDAC organoids with recent data from primary PDAC ^19^. Across all patients, PDAC organoid cells were uniformly identified as malignant ductal cells by reciprocal principal component analysis and expression of characteristic genes (Supplementary Figure 1B), consistent with previous observations that the *in vitro* culture conditions promote ductal cell growth ^21^.

### PDAC organoids show patient-specific expression of distinct gene families

PDAC is characterised by a small number of recurrent genetic alterations that occur at high frequency, and therefore cannot account fully for differences in disease progression and therapeutic response between patients ^9^. To investigate distinguishing features of PDAC organoids at the transcriptional level, we constructed a shared nearest neighbour (SNN) graph of all cells and found that cells from the same patient clustered together (Figure 1C), even if they derived from different biopsies or metastatic sites processed separately. This indicates that transcriptional differences between patients are larger than within patients and are not merely due to technical batch effects. Patient-specific differences in transcriptional profiles were also not explained by individual known expression quantitative trait loci ^24^ (Supplementary Figure 1C).

To further investigate the origin of transcriptional heterogeneity between PDAC tumors, we determined genes that were differentially expressed in organoid models of primary tumor cells derived from each patient compared to all others. While expression of the most highly upregulated genes in each PDAC organoid was often patient-specific (Figure 1D,E and Supplementary Table 2), we observed that many of these genes belonged to the same gene families. Analysis of gene family membership of the top 20 enriched genes for each patient, excluding gene families defined by specific molecular domains, showed that the most highly represented gene families included cell surface and transmembrane proteins (CD molecules, claudins, solute carriers), secreted proteins that interact with the extracellular matrix (mucins, kallikreins), enzymes that metabolise a range of substrates and potential drug targets (serine proteases, aldo-keto reductases), and type-I keratins (Figure 1F). In PDAC organoids where these genes were detected, they were frequently expressed in a large proportion of cells, but with very low or zero expression in other lines (Figure 1G). Since many of the gene families we identified have previously been proposed as indicators of tumor identity and/or prognosis in PDAC patients ^25–30^, these results have important implications for biomarker identification in PDAC, highlighting that samples from a large enough cohort of patients need to be analysed in order to overcome interpatient transcriptional heterogeneity.

### Transcriptional subtype heterogeneity correlates with poor prognosis

PDAC is commonly classified into ‘classical’ and ‘basal-like’ transcriptional subtypes, with the latter carrying a poorer prognosis ^14^. Bulk RNA-seq of PDAC organoids from our cohort grouped organoid lines into basal-like, classical and intermediate subtypes (Supplementary Table 1). To elucidate subtypes at the single-cell level, cells from all patient-derived organoids were scored for the expression of published subtype signatures consisting of 25 genes per subtype ^14^. Overall, subtype annotation correlated well with bulk results (Supplementary Table 3). For the majority of patient-derived organoids, following PCA-based clustering of cells, subtype signature scores were homogeneous across cell clusters and reflected the subtype determined by bulk RNA-seq (Figure 2A). Conversely, we identified a subset of patient-derived organoids containing both ‘basal-like’ and ‘classical’ cells (Figure 2B), with corresponding marker gene expression (Figure 2C). Notably, we found that PDAC organoids exhibiting heterogeneous subtype identity had been classified as WHO grade 3 or 4 tumors in histopathological assessments, whereas PDAC organoids with homogeneous subtype identity had been classified as grade 2 or 3 (Figure 2D). Moreover, patients in our cohort with homogeneous classical PDAC tended to have longer overall survival whereas the outcome of basal-like tumors tended to be poorest (Supplementary Figure 1D). Homogeneous enrichment for ‘classical’ subtype marker genes in PDAC organoid lines therefore correlates with more differentiated tumors, which are associated with a better prognosis.

**Figure 2:**
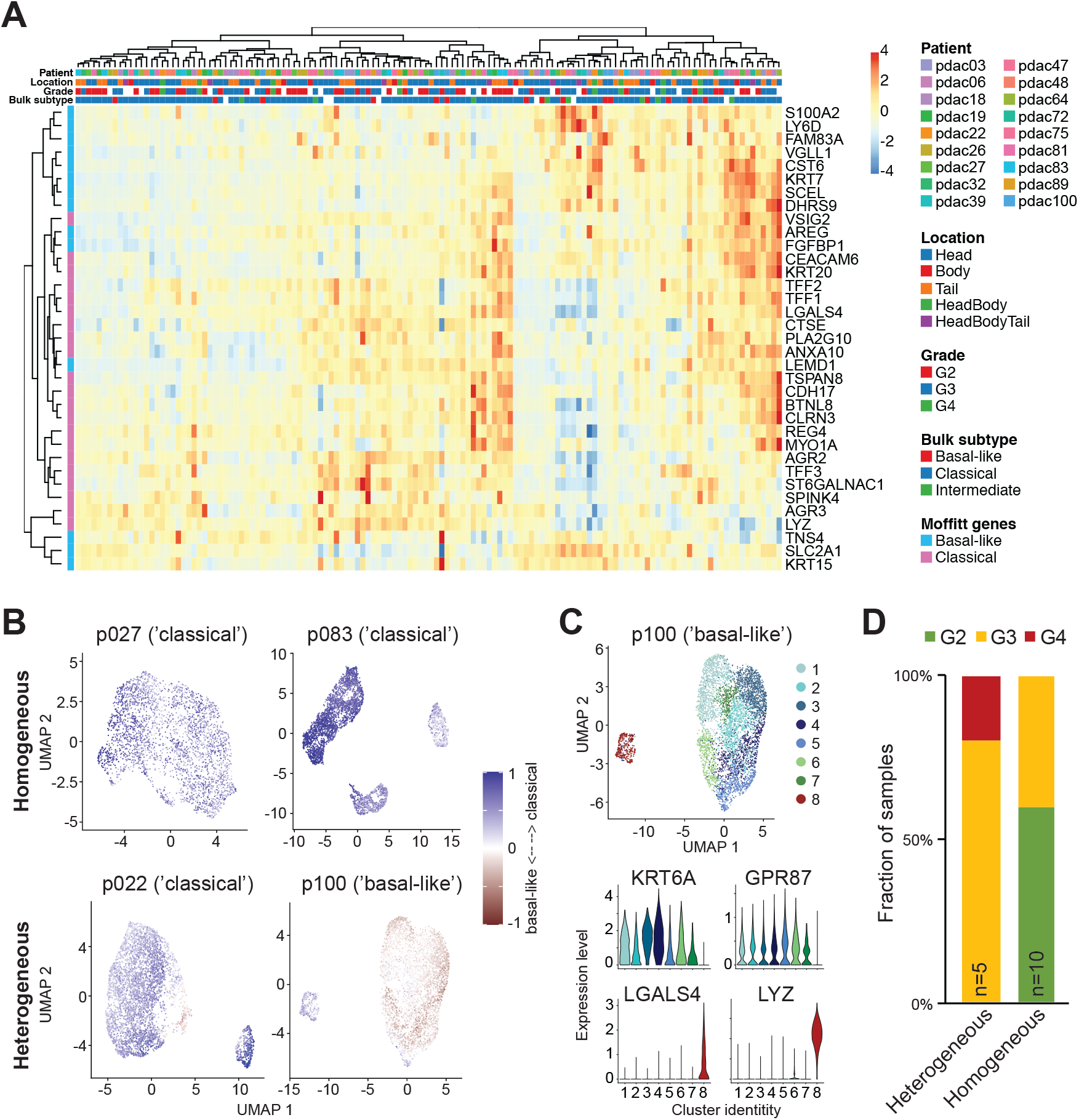
Expression of PDAC subtype signatures. **A)** Heatmap showing the expression of Moffitt subtype signature genes across individual clusters from all patients. **B)** Distribution of Moffitt subtype scores across PDAC organoid cells, shown for two homogeneous (top) and two heterogeneous (bottom) samples. Blue indicates classical subtype scores, red basal-like (Methods). **C)** Top: Eight distinct clusters of cells were identified in p100. Bottom: Expression of characteristic genes for the basal-like (top) and classical (bottom) subtype across clusters, showing significantly higher expression of classical genes and lower expression of basal-like genes in cluster 8 compared to all others. **D)** Proportion of homogeneous and heterogeneous organoids that were classified as grade 2, 3 or 4 by histopathological assessment of the original tumor.

### Functional cell states are shared across patients

To identify cellular states that are shared across tumors, we performed reciprocal PCA-based integration of 18 primary PDAC organoid transcriptomes (Methods). Clustering of cells revealed nine functional cell states that were conserved across patients (Figure 3A and Supplementary Figure 2A).

**Figure 3:**
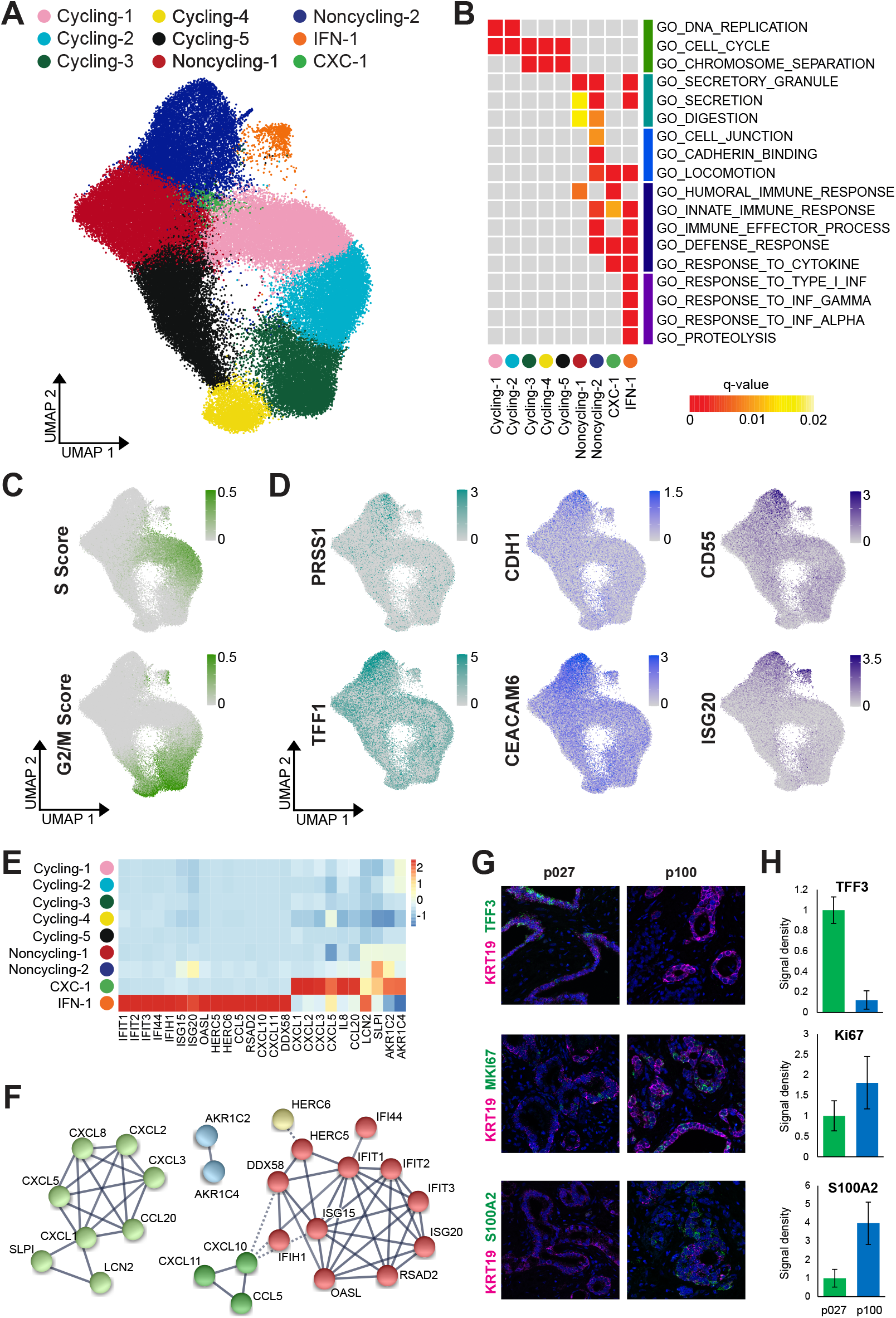
Functional cell states shared across PDAC patients. **A)** UMAP representation of clusters shared across patients, after PCA-based integration of PDAC transcriptomes from 18 primary PDAC organoid lines. **B)** Gene ontology (GO) term enrichment analysis using the top 100 upregulated genes per cluster. Grey boxes indicate no significant enrichment. **C)** Cell cycle scores for each cell, computed based on the expression of S and G2/M phase genes, were visualized using the same UMAP representation as in (A). **D)** Expression of characteristic genes across all cells from primary PDAC organoid lines, visualized using the same UMAP representation as in (A). **E)** The heatmap shows the average expression per cluster of genes specifically enriched in clusters CXC-1 and IFN-1 with interactions recorded in the STRING database ^52^. **F)** Networks representing gene interactions considered in (E). **G)** Representative images of RNA fluorescence in situ hybridization (FISH) staining for KRT19 indicating PDAC cells (magenta) together with either TFF3, MKI67 or S100A2 (green) in primary tumour sections from p027 and p100. Nuclei are stained with DAPI (blue). Scale bar, 50 µm. **H)** Quantification of RNA FISH stainings shows the signal density per nucleus for each transcript in PDAC cells (identified by KRT19 staining), normalized to the density in sample p027 (p027: n=339 nuclei for MKI67 and TFF3, 369 nuclei for S100A2; p100: n=176 nuclei for MKI67 and TFF3, 306 nuclei for S100A2; each from 3 separate images, with error bars showing standard errors in the mean).

We evaluated the characteristics of each cluster by differential expression analysis (Figure 3B and Supplementary Table 4). Five clusters represented cell moving through the different cell cycle phases (labelled Cycling-1 to Cycling-5). Two clusters were devoid of cycling cells (labelled Nonycling-1 and Noncycling-2), but comprised cells expressing the cyclin dependent kinase inhibitor 1 (Supplementary Figure 3A). These clusters showed increased expression of genes involved in secretion, digestion, cell adhesion and locomotion. While immune response signalling was generally upregulated across the noncycling clusters, one additional cluster (labelled IFN-1) showed specific expression of genes involved in type I interferon signalling. The smallest cluster (labelled CXC-1) was enriched for cells expressing CXC motif ligands such as *CXCL1*, *CXCL2* and *CXCL8*, which are thought to stimulate cancer cell proliferation and migration ^31–33^. Clusters IFN-1 and CXC-1 both comprised cycling and quiescent cells (Supplementary Figure 3B). Notably, the Cycling and Noncycling clusters contained cells from all patient samples, but only five patients (p006, p018, p027, p047 and p089) contributed at least 1% of sampled cells to cluster IFN-1 and only two patients (p064 and p100) contributed similarly to cluster CXC-1 (Supplementary Table 5). RNA in situ hybridization confirmed the differential representation of clusters IFN-1 and CXC-1 in patient-derived cultures as well as primary tumors (Supplementary Figure 2B).

We thus concluded that all patient-derived PDAC organoids contained cycling cells, which we resolved into different cell cycle phases (Figure 3C), and differentiating cells assuming functional characteristics reminiscent of their pancreatic origin (Figure 3D). In addition, a subset of patients also harbored cell clusters with specific expression of cytokines thought to contribute to tumor progression by autocrine and paracrine mechanisms (Figure 3E,F).

RNA in situ hybridization of surgical primary tumor samples for KRT19 (a marker of neoplastic PDAC cells), TFF3 (a marker of secretory cells and of the classical PDAC subtype), S100A2 (a marker of the basal-like PDAC subtype) and MKI67 (a proliferation marker) showed higher TFF3 expression in the classical subtype and higher S100A2 expression in the basal-like subtype (Figure 3G,H), confirming that PDAC organoids reflect the original patient tumors.

### Differentiation hierarchy in PDAC organoids

Based on the observation that a subset of clusters shared across PDAC organoid lines contained cycling cells, we sought to identify the differentiation trajectory of PDAC organoid cells. Applying the recent concept of RNA velocity ^34^, we determined changes over time in gene expression, as estimated by the relative abundance of unspliced and spliced mRNA in each sample (Figure 4A). By Markov chain tracing, we identified probable startpoints and endpoints of differentiation trajectories (Figure 4B). Across all patients, trajectory startpoints coincided with cycling clusters, with most trajectories converging onto cluster Cycling-4 as the cell population of origin (Figure 4B,C). Trajectory endpoints, on the other hand, were predominantly located in the distant cluster Noncycling-2 (Figure 4B,C).

**Figure 4:**
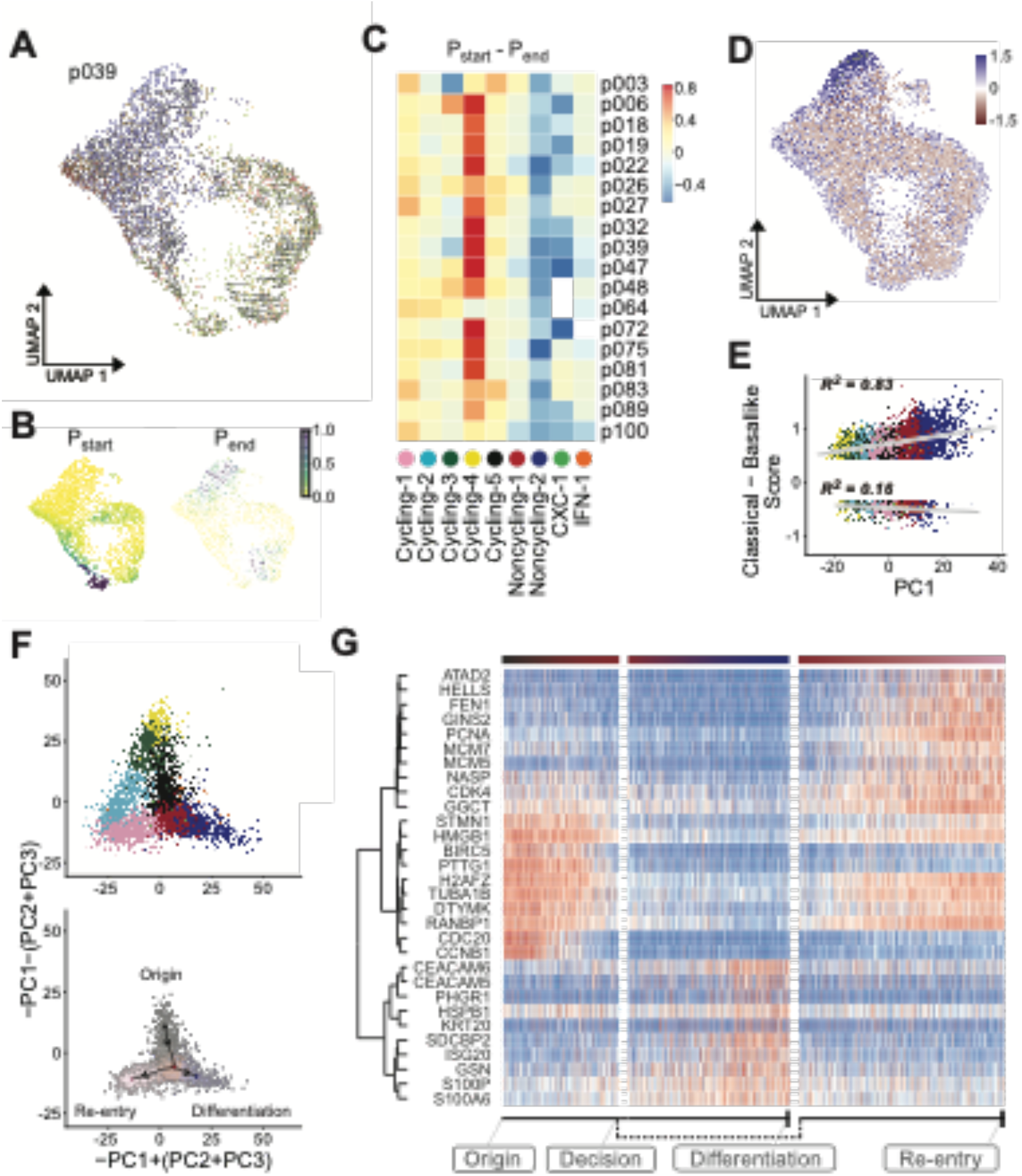
Differentiation hierarchy of PDAC cell states *in vitro* and *in vivo*. **A)** RNA velocity trajectory shown for p039, using the same UMAP representation as in Figure 3A. **B)** Startpoint (top) and endpoint (bottom) probabilities shown for p039. **C)** Difference between the average startpoint (red) and endpoint (blue) probabilities within each cluster (columns) across all PDAC organoid samples (rows). **D)** Distribution of PDAC subtypes in single cells along the lineage trajectory. Colors indicate whether a cell exhibits higher expression of ‘classical’ (blue) or ‘basal-like’ (red) subtype marker genes. **E)** Correlation of PDAC subtype scores with cell embeddings in the first principal component, which distinguishes cycling cells (negative values in PC1) from differentiating cells (positive values in PC1). **F)** Projection of all cells onto a linear combination of PC1, PC2 and PC3 to visualize cycling and differentiating cells (top). Cells in the four clusters around the trajectory bifurcation point were linked by a minimum spanning tree (bottom). **G)** Expression of genes with dynamic changes along the bifurcation branches (red: high, blue: low), with hierarchical clustering of gene expression shown on the left. Column annotations indicate position along the branches.

Notably, when relating the Moffitt subtype identity of cells to their position along the lineage trajectory, we found that the ‘classical’ gene signature was enriched in the differentiated secretory compared to the cycling cells (Figure 4D,E).

As our *in vitro* organoid model is enriched for progenitor-like cells compared to primary tumor samples (Supplementary Figure 1B), different cell cycle phases could be resolved in our data, and we identified a bifurcation point in G1 phase at which cells either re-enter the cell cycle or differentiate towards the secretory state (Figure 4F). To further investigate the fate behaviour of PDAC organoid cells, we calculated a minimum spanning tree linking all cells in clusters Cycling-1, Cycling-5, Noncycling-1 and Noncycling-2, and assigned each cell a pseudotime value along the bifurcating path. In addition to ubiquitous cell cycle related genes, this approach identified potential drivers of cellular fate; for example, *GGCT* (gamma-glutamylcyclotransferase) and *RANBP1* (Ran binding protein 1) show increased expression soon after the bifurcation point in cells re-entering the cell cycle, whereas the cell adhesion gene *CEACAM6* and *S100A6* encoding a calcium-binding protein are upregulated in differentiating cells (Figure 4G).

We thus conclude that PDAC organoid growth is fuelled by a large pool of cycling cells, which sustain a population of differentiating progeny that assume secretory function. Notably, this functional hierarchy is conserved in PDAC organoids from all patients in our cohort, as well as in scRNA-seq data from primary PDAC (Supplementary Figure 2C).

### Liver metastases re-establish functional aspects of the primary tumour

PDAC is frequently metastatic at diagnosis, with only 10-20% of patients qualifying for a potentially curative surgery ^35^. While most patients with advanced PDAC receive palliative chemotherapy, the benefit of neoadjuvant chemotherapy is a subject of current debate ^36, 37^. A recent bulk whole-genome sequencing study of treatment-naïve PDAC tumors and metastatic lesions found identical known driver gene mutations in primary tumors and all matched metastases ^12^, spurring hope that targeted therapies could therefore provide a clinical benefit in advanced PDAC by simultaneously eliminating genetically homogeneous cells at multiple sites.

Here, we addressed the question of cellular heterogeneity in metastatic PDAC at the transcriptional level, by deriving PDAC organoid lines from two liver metastases (M1 and M2) as well as from the primary tumor from the same patient (P). Following scRNA-seq and merging the three transcriptomes by reciprocal PCA, we identified five cell clusters with differential representation in the three PDAC organoid lines (Figure 5A). All samples contained cycling cells, as indicated by *MKI67* expression (Figure 5B). Other functional aspects of PDAC, including secretion (e.g. *FABP1*, *SCG5*), digestive enzymes (e.g. *PRSS1*, *PGC*), cytoskeleton or cell adhesion genes (e.g. *ANXA6, TUBB2B*, *CEACAM6*), MHC complex members (e.g. *CD74*, *HLA-DQB1*, *HLA-DRB1*) and putative inflammasome inhibitory genes (e.g. *TMEM176A, TMEM176B)* ^38^, were heterogeneously expressed between clusters (Figure 5B). For example, cluster 2, which was enriched for genes related to secretion and digestion, was detected in P and M2, but not in M1.

**Figure 5:**
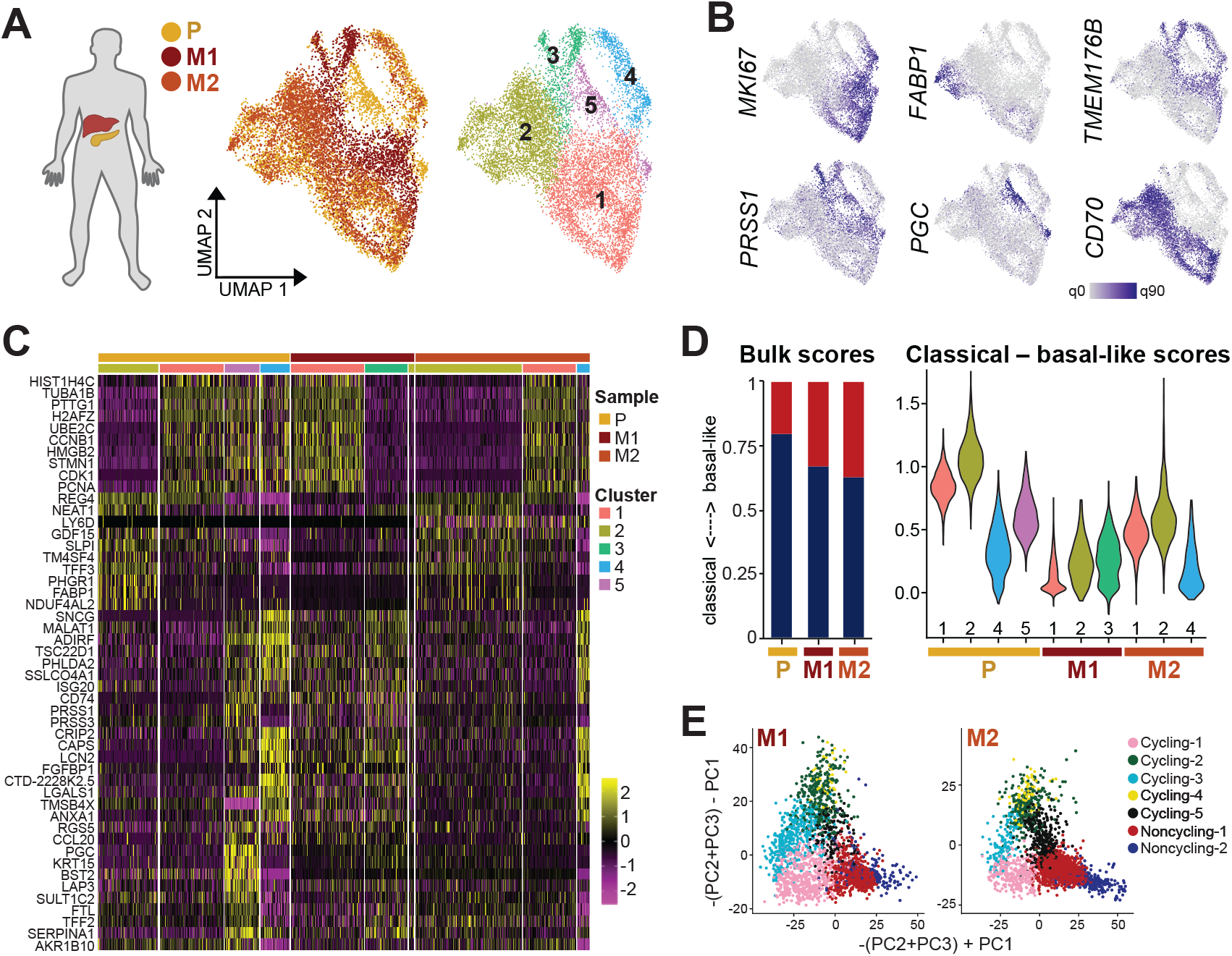
Functional and transcriptional subtype heterogeneity in liver metastases. **A)** For one patient, organoid lines were derived from the primary tumor (P) as well as two liver metastases (M1 and M2). Shown is a UMAP representation of transcriptomes integrated by reciprocal PCA, with shared and organoid line-specific cell clusters identified by unsupervised clustering. Cells are colored by sample origin (left) and cluster membership (right). **B)** Expression of genes involved in cell cycle (*MKI67*), secretion and digestion (*FABP1*, *PRSS1, PGC*), and immune regulation (*CD70*, *TMEM176B*) across cells from P, M1 and M2 organoids. **C)** Expression of the top 10 enriched genes for each cluster. **D)** Moffitt subtype scores from bulk RNA-seq data (left) and single-cell RNA-seq data (right) for P, M1 and M2 organoids. Blue indicates classical subtype scores, red basal-like (Methods). Cells are grouped by cluster identity according to (A) in the violin plots. **E)** Projection of M1 and M2 transcriptomes onto the primary PDAC organoid data (see Figure 4F and Methods) shows distribution of cells across cycling and differentiating clusters.

Bulk RNA-seq suggested that both metastasis-derived organoid lines showed more ‘basal-like’ gene expression than organoid lines derived from the primary tumor (Supplementary Table 1), consistent with earlier observations of ‘basal-like’ phenotype enrichment in metastatic tissues^14^. Conversely, at the single cell level, we found transcriptional subtype heterogeneity even within a single metastasis-derived organoid line, with a subset of cells from M2 showing more ‘classical’ gene expression than a subset of cells of organoids derived from the primary tumor (Figure 5D).

By projecting scRNA-seq data from both metastasis-derived organoid lines onto the PCA of all primary PDAC organoid transcriptomes, we further observed a depletion of differentiated cells (cluster ‘Noncyling-2’) especially in M1 (Figure 5E), corresponding to the depletion of functional clusters 2 and 3 identified above (Supplementary Figure 3C).

These observations suggest that metastatic PDAC lesions in the liver re-build diverse aspects of the original tumor, including the differentiation hierarchy identified in primary samples. Since both M1 and M2 express genes also detected in the primary tumor but not shared between the metastases, the observed gene expression pattern is consistent with an underlying homogeneous mutational landscape in combination with differential epigenetic control, possibly driven by the local microenvironment, or chance events. While PDAC metastases thus appear more heterogeneous at the transcriptional compared to the genomic level ^12^, differences in gene expression may not be the result of *de novo* events, with encouraging implications for future targeted therapies.

### PDAC organoids provide an *in vitro* model for imaging-based drug screens

In the quest to develop novel chemotherapies for PDAC patients with advanced metastatic disease or unfavourable transcriptional subtypes, organoid models offer great potential for *in vitro* drug response screens. To illustrate the utility of our PDAC organoid lines for screening applications, we applied a recent automated microscopy-based live cell assay and quantification workflow (*DeathPro*) ^39^. In this workflow, all cells are stained with Hoechst 33342 and propidium iodide (PI) to distinguish live and dead cells; the total areas covered by dead cells (PI) and all cells (Hoechst 33342 or PI stained) are then measured from projected confocal images and used to calculate area-under-curve values for cell death (AUCd) and proliferation inhibition (AUCpi). Compared to simpler luminescent cell viability assays, this approach resolves drug-induced cell death and proliferation inhibition ^39^. Here, we measured PDAC organoid cell death and proliferation inhibition induced by six drugs in clinical use for PDAC therapy; these were 5-FU, Gemcitabine, Irinotecan, Paclitaxel, Erlotinib and Oxaliplatin (Figure 6A and Supplementary Table 6).

**Figure 6:**
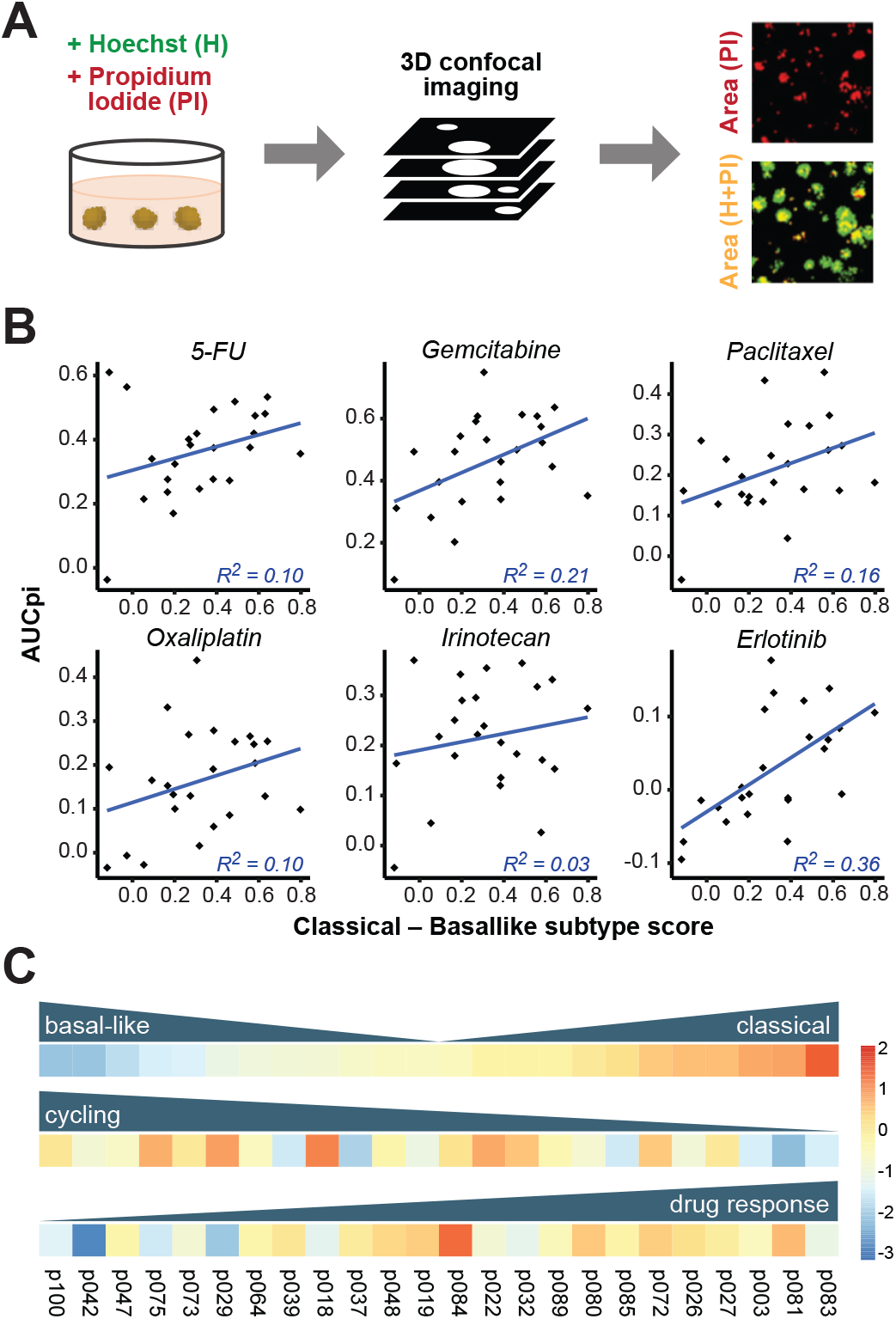
Classical subtype gene expression correlates with organoid drug responses. **A)** Overview of the experimental drug screen protocol. PDAC organoids were treated with a range of drug concentrations over 72 hours, and then subjected to 3D confocal imaging. By measuring the intensity of Hoechst (H) and propidium iodide (PI) signals, scores were computed to compare drug-induced cell death and proliferation inhibition. Scale bars, 200 µm. **B)** Correlation between median subtype scores calculated from scRNA-seq data and drug-induced proliferation inhibition, as measured by the area under curve (AUCpi), for 24 PDAC organoid lines. **C)** Overview of variation in transcriptomic PDAC subtype (top), median cell cycle score (middle) and proliferation inhibition in response to 5-FU treatment (bottom), shown as z-scores scaled across all samples.

5-FU and Gemcitabine elicited the most cell death and proliferation inhibition across all organoid lines included in the screen, with Erlotinib and Oxaliplatin the least effective, although significant heterogeneity in drug responses was observed between organoid lines (Supplementary Figure 4A). For each drug, ‘classical’ subtype gene expression was correlated with more pronounced proliferation inhibition in response to drug treatment, whereas drug-induced cell death showed less correlation with PDAC subtype (Figure 6B and Supplementary Figure 4B).

Taken together, our results indicate that the ‘classical’ subtype signature is linked to better drug responses and a higher level of differentiation, while the ‘basal-like’ signature with poorer prognosis is linked to cell proliferation and reduced drug responses (Figure 6C and Supplementary Figure 4C).

## Discussion

PDAC remains a challenging entity for experimental study; *ex vivo* tumor samples are difficult to access and often have low malignant cell content ^15^. Recent advances in *in vitro* organoid technologies have therefore sparked hope for an alternative route. We have shown here that PDAC organoids established with our protocol are transcriptionally equivalent to malignant ductal cells. Notably, our results strongly overlap with previously published primary PDAC scRNA-seq data ^19^, suggesting that PDAC organoid cells recapitulate key aspects of cell state heterogeneity in the malignant ductal cell compartment *in vitro*.

By deriving primary PDAC organoids from 18 patients, we found that PDAC organoids show patient-specific differences that mask the shared cell state heterogeneity we also identified. This observation highlights the need to include a sufficiently large number of patient samples in single-cell studies in order to distinguish common features from individual variation, enabling extrapolation to a larger patient collective. We observed frequent differential expression of genes encoding cell surface and secreted proteins between patients, often from the same gene families (e.g. claudins, mucins). These transcriptional differences may thus not always have functional relevance, but could reflect underlying genetic or epigenetic heterogeneity, different cells of origin or microenvironments of the tumor, lifestyle and comorbidities of the patient, or differences in therapy and immune response. It will be interesting to explore the origin of patient-specific transcriptome profiles in future studies and determine if this heterogeneity is also present in healthy pancreas cells or arises during tumorigenesis.

To categorize transcriptional heterogeneity in PDAC, an established classification scheme distinguishes ‘classical’ from ‘basal-like’ tumors. In line with recent results from other groups ^17, 40^, our cohort comprised PDAC organoids with uniformly ‘classical’ gene expression, but also a subset of organoids with heterogeneous expression of subtype marker genes across cells. The latter were associated with a poorer prognosis of the patient, as determined by histopathological grading of the original tumor and patient outcomes. As our cohort did not include any tumors that were purely ‘basal-like’, it was impossible to resolve whether poorer prognosis simply results from higher ‘basal-like’ cell content or whether the extent of subtype heterogeneity is an independent predictor, which thus remains an interesting question for future studies. It should also be noted that the relative paucity of basal-like cells in our PDAC organoids may reflect selective pressure exerted by the *in vitro* culture conditions towards ‘classical’ subtype gene expression. Nevertheless, the broad agreement of organoid subtype assignments with original tumor subtypes indicates that differences between PDAC samples are retained in culture, consistent with an earlier study in which PDAC organoid drug responses paralleled patient outcomes ^41^.

Despite substantial interpatient heterogeneity, our results identified functional cell states that are shared across patients, and are linked by a differentiation hierarchy conserved *in vitro* and *in vivo*. Proliferating cells may either re-enter the cell cycle, or give rise to differentiating cells which acquire characteristics of pancreatic tissue including digestive and secretory functions. Expression of genes involved in digestion and secretion has previously also been observed in primary PDAC, again confirming that our organoid model captures key aspects of this entity ^42^. While our analyses provide initial insight into differential gene expression at the bifurcation point, future studies should address whether the fate choice of tumor cells could be exogenously biased towards differentiation in order to constrain tumor growth. In addition, it will be interesting to investigate whether a subset of cells in clusters ‘Noncycling-1’ and ‘Noncycling-2’ may be able to re-enter the cell cycle, which would enhance the growth potential of the tumor.

Importantly, malignant cells may express a combination of transcriptional programmes that are not normally found within the same cell type in the healthy pancreas. By comparing scRNA-seq data from organoid lines derived from two liver metastases to the matched primary tumor, we also found that metastases may activate different combinations of pancreatic transcriptional programmes. As PDAC metastases are known to share the same driver mutations as the primary tumor ^12^, heterogeneity in PDAC metastases therefore most likely arises from differential epigenetic regulation activating or deactivating transcriptional programmes.

The expression of pancreatic programmes in liver metastases of PDAC also resolves a debate regarding the role of normal pancreas genes in PDAC transcriptome data. Due to the frequently low tumor cellularity of PDAC, it has been questioned whether genes that are normally expressed in mature pancreatic tissue should be excluded from analyses of PDAC pathophysiology, assuming that they represent normal tissue contamination ^18^. Not surprisingly, a recent study found an association between the presence of normal pancreatic transcripts and the presence of healthy tissue contamination on PDAC tumor slides ^43^. Our results indicate that such genes may in fact be expressed by malignant cells in PDAC tumors, even in distant metastatic lesions, and their inclusion in future studies of PDAC pathophysiology is therefore warranted.

In addition to proliferating cells and differentiating cells with pancreatic expression programmes, a subset of PDAC organoid lines contained distinct clusters of cells that may mediate type I interferon signalling (cluster IFN-1) or CXC motif cytokine signalling (cluster CXC-1). As these signalling pathways have been implicated in cancer progression and metastasis as well as therapy responses ^31–33, 44^, resolving the presence of corresponding cell populations by single-cell transcriptome studies could improve prognostic accuracy for patients in the future.

Our drug screen showed that drug response, both at the level of cell death and proliferation inhibition, was positively correlated with the expression of ‘classical’ subtype genes, and negatively correlated with ‘basal-like’ gene expression. PDAC organoid lines with higher drug response also tended to display a more differentiated transcriptome, whereas lower drug response was associated with higher cell cycle scores. These results are consistent with ‘basal-like’ PDAC tumors showing poorer response to standard therapy and worse prognosis in the clinic. Our organoid-based assay thus represents a promising approach for the *in vitro* identification of compounds that could improve therapeutic outcomes for PDAC patients.

## Materials and Methods

### PDAC organoid generation and maintenance

Pancreatic tumor organoid cultures were established using an adaptation of the previously described protocol ^21^. Pancreatic tumor specimens were obtained from patients undergoing surgical resection at the Surgical Department of the Heidelberg University Hospital. Written informed consent from all patients was obtained prior to acquisition of tissue. Tumor specimens were minced and digested for up to 4 h at 37°C in AdMEM/F12 medium (Gibco) containing 2 mM GlutaMAX (Gibco), 10 mM HEPES (Gibco), 1x Primocin (InvivoGen)), 1 mg/ml Collagenase type IV (Sigma-Aldrich), 100 µg/ml DNase I (AppliChem), 1x B27 (Gibco), 1 mM N-acetylcysteine (Sigma-Aldrich) and 10 µM Y-27632 (Selleckchem). Dissociated cells were seeded in Growth Factor Reduced Matrigel (Corning) and cultured in growth organoid medium consisting of AdMEM/F12 medium, 2 mM GlutaMAX, 10 mM HEPES, 1x Primocin, 1x B27, 1 mM N-acetylcysteine, 10% RSPO1-conditioned medium ^45^, 100 ng/ml FGF10 (PeproTech), 100 ng/ml Noggin (PeproTech), 500 nM A83-01 (Tocris) and 10 µM Y-27632. Medium was refreshed every 3-4 days omitting Y-27632. Organoids were routinely passaged by dissociation with TrypLE (Gibco) for 10 min at 37°C. Medium was further supplemented with 50 ng/ml EGF (PeproTech) and, if required, with 50% Wnt3A-conditioned medium ^46^ only after tumor cell enrichment to avoid overgrowth of normal ductal cells during the initial passages. PDAC organoids were cultured for 2.2 months on average before bulk RNA-seq (range: 1.0 - 4.7 months), and 3.6 months on average before scRNA-seq (range: 1.9 - 5.8 months). The technical replicates p039 and p039b were cultured for 2.8 months and 4.2 months, respectively, before scRNA-seq.

### Single-cell dissociation, library preparation and sequencing

Single-cell sequencing libraries were prepared according to the 10x Genomics Single Cell 3 v2 Reagent Kit user guide with small modifications. Organoid cultures were expanded in growth medium without addition of Wnt3A-conditioned medium (serum-free condition) for 5 days and dissociated into single cells for 30-60 min using AccuMax (Invitrogen) supplemented with 0.3 mg/ml Dnase I. Cell were washed with PBS containing 0.04% BSA, strained through a 20 µm strainer (PluriSelect) and counted. Single-cell suspensions containing 10,000 cells were loaded following the protocol of the Chromium Single Cell 3’ Library Kit v2 (10x Genomics) to generate cell and gel bead emulsions. After droplet generation, samples were transferred onto a pre-chilled 96-well plate (Eppendorf), heat-sealed and reverse transcription was performed using a Bio-Rad C1000 Thermal Cycler. After reverse transcription, cDNA was recovered using Recovery Agent followed by Silane DynaBead clean-up. Purified cDNA was amplified for 15 cycles and cleaned up using SPRIselect beads (Beckman). Samples were quantified on an Invitrogen Qubit 4 Fluorometer. cDNA libraries were prepared according to the Single Cell 3 Reagent Kits v2 guide with appropriate choice of PCR cycle number based on the calculated cDNA concentration. Final libraries were sequenced in one lane per sample with the Illumina NextSeq 500 system in high-output mode (paired-end, 75 bp).

### scRNA-seq analysis

#### Alignment and quality control

Raw sequencing data were processed using CellRanger version 2.1.1 (10x Genomics). Transcripts were aligned with the 10x reference human genome hg19 1.2.0. Quality control and downstream analysis were performed with Seurat version 3.0 ^47^. Cells with fewer than 200 genes or genes represented in fewer than 3 cells were excluded from the analysis. Cells with >100.000 reads or >15% mitochondrial reads were excluded. Count data was log-normalized with a scale factor of 10.000, and the 2000 most variable genes identified with the FindVariableFeatures function in Seurat. Normalized data were scaled using the ScaleData function.

#### Analysis of individual organoid samples

For the analysis of individual samples, organoid scRNA-seq data from primary tumors and metastases was processed with the Seurat package in R, using the FindClusters function with a resolution of 0.5. To identify differences between patients, expression data from all samples was combined and clustered using the FindClusters function with a resolution of 0.1, resulting in one cluster per organoid except for pdac100, for which we manually combined two resulting clusters. Dimensional reduction was performed using the uwot package. Differentially expressed genes for primary PDAC samples were identified using a Wilcoxon rank sum test. As p080 and p081 were derived from the same patient, only the latter was included in this analysis. The top 20 enriched genes per sample that were expressed in at least 25% of cells in this sample and at most 90% of cells in all others, were analysed according to their gene group membership.

#### Moffitt subtype analysis

To analyse Moffitt subtypes, cells from each patient were clustered using the FindClusters function in Seurat with a resolution of 0.5. The average expression of 25 marker genes each for the ‘classical’ and ‘basal-like’ subtype ^14^ was then determined for each cluster. The gene expression heatmap was generated using complete clustering of Spearman correlation coefficients. Subtype scores for basal-like (*S_bas_*) and classical (*S_cla_*) marker gene expression each individual cell were calculated using the AddModuleScore function in Seurat, and a combined score (*S_moff_* = max[0, *S_cla_*] – max[0, *S_bas_*]) was calculated for each cell.

#### Joint analysis of PDAC samples by reciprocal PCA

To analyse cell states across patients, primary tumor samples were merged by reciprocal PCA. As p080 and p081 were derived from the same patient, only the latter was included in this analysis. Cells were clustered in Seurat using the FindClusters function with the Louvain algorithm with a resolution of 0.7, and dimensional reduction was performed using the umap-learn package ^48^. Differentially expressed genes for each cluster were identified using a Wilcoxon rank sum test. Gene ontology (GO) term analysis was performed using the MSigDB ^49, 50^ with the top 100 enriched genes per cluster with adjusted p-value <0.05. One cluster resulted in only six enriched genes and no enriched GO terms but comprised cells with the highest number of RNA counts in the dataset, suggesting a technical artefact; it was therefore excluded from the analysis. Cell cycle scores were calculated with the AddModuleScore function in Seurat, using previously published lists of S phase and G2/M phase marker genes^51^. We used the STRING database ^52^ to identify gene interaction networks specifically expressed in clusters, with the confidence cut-off at 0.75 and k-means clustering with k = 5.

#### Comparison to primary PDAC scRNA-seq data

To compare scRNA-seq data from our PDAC organoids with primary PDAC, we downloaded raw data for two patients (T18 and T20) from a recent publication ^19^. Primary PDAC data was processed analogously to PDAC organoid data, and dataset integration was performed using reciprocal PCA in Seurat, with primary PDAC as the reference. Cell type identification was based on established marker genes ^19^.

#### Inference of a lineage hierarchy

We used the velocyto Python package ^34^ to estimate RNA velocities by distinguishing unspliced and spliced mRNAs. Correlations of *S_moff_* with PC1, which resolved cycling from differentiated cells, were calculated by linear regression using mean values across cells within bins of equal width in both *S_moff_* and PC1. For visualization, only cells with greater than median *S_moff_* for *S_mof_* > 0 or smaller than median *S_moff_* for *S_mof_* < 0 were plotted. Linear combinations of cell embeddings for the first three principal components, specifically –PC1+(PC2+PC3) and –PC1–(PC2+PC3), were chosen to illustrate the branching point at which cells either re-enter the cell cycle or differentiate. Following this two-dimensional projection, a randomly sampled subset of 3,000 cells in clusters Cycling-1, Cycling-5, Noncycling-1 and Noncycling-2 were located along a bifurcating trajectory by a minimum spanning tree. Expression of genes varying along the trajectory was visualized using the dynverse package in R ^53^. To determine if an equivalent trajectory could be resolved *in vivo*, two primary PDAC datasets (T18 and T20) from a recent publication ^19^ were processed analogously.

#### Comparison of metastases to the primary tumor

To compare the transcriptome of PDAC organoids derived from two metastases (p084 and p085) to organoids derived from the corresponding primary tumor (p083), transcriptomes from these three samples were merged by reciprocal PCA, and cells were clustered with a resolution of 0.3. Expression of the top 10 upregulated genes for each cluster was visualised in a heatmap. Subtype scores were calculated as described above. For trajectory analysis, integrated gene expression data was projected onto the PCA of all primary tumor samples, using the first 50 principal components. Cluster identity of metastases-derived cells was determined according to their similarity to primary tumor-derived cells, using the TransferData function in Seurat.

### RNA Fluorescence In Situ Hybridization

Sections of PDAC organoids or primary tumor samples were processed for RNA in situ detection using the RNAscope Multiplex Fluorescent Reagent Kit v2 according to the manufacturer’s instructions (Advanced Cell Diagnostics). RNA fluorescence in situ hybridization (FISH) images were acquired on a Leica SP8 confocal laser scanning microscope equipped with a 40×/1.30 oil objective (Leica HC APO CS2). Images were binarized, Gaussian filtering followed by watershed segmentation was applied to identify nuclei, and binarized FISH signal density per nucleus was calculated using ImageJ.

### Imaging-based drug screen

The organoid-based drug screen was performed using the ‘DeathPro’ workflow as previously described ^39^ on a subset of 24 PDAC organoid lines, including one derived from an unmatched peritoneal metastasis (p073) and one derived from an unmatched perivascular metastasis (p037). Briefly, drugs were dissolved in DMSO, water, PBS or ethanol as required sand stored as single-use aliquots at −80°C. Drug dilution series (1:3) were prepared in PDAC organoid culture medium containing 1 μg/ml Hoechst (Invitrogen) and 1 μg/ml PI (Sigma). Organoids were incubated in Matrigel-coated 96-well Angiogenesis μ-Plates (ibidi) with drug-containing medium for 72 h. Organoid cultures were then washed twice with PBS and the medium was exchanged for drug-free medium. Cells were imaged 72 h after the start of drug treatment using a Zeiss LSM780 confocal microscope, 10× objective (EC Plan-Neofluar 10×/0.30 M27) and 405 and 561 nm diode lasers in simultaneous mode. Imaging was performed in an incubation chamber at 37°C, 5% CO_2_ and 50-60% humidity using the Visual Basic for Applications macro ‘AutofocusScreen’ ^54^. Binarized images from Hoechst and PI channels were combined (H + PI) to calculate total cell area, while PI alone was used as a proxy for dead cells. Drug response curves to determine the LD50 were fitted if there was a significant difference between cell death in drug-treated and untreated samples (ANOVA with *P*-values < 0.0005), and area under curve (AUC) values were calculated for cell death. Cell growth was estimated by dividing total cell area (H + PI) at 72 h by total cell area (H + PI) at 0 h after drug application, and the area under curve for proliferation inhibition (AUCpi) was determined. To correlate drug responses with scRNA-seq results, median cell cycle scores were calculated for each sample with the AddModuleScore function in Seurat, using previously published cell cycle genes ^51^.

### Comparison of overall survival times

Overall survival (OS) was defined as the time from surgery to death or last follow-up. OS shorter than 90 days after surgery was considered peri-operative death and therefore excluded from the survival analyses. Median survival was estimated using the Kaplan–Meier method, patients still alive at the last follow-up were censored. Survival curves between groups were compared by the log-rank test. Analysis and plots were performed using the R packages survival and survminer.

## Data availability

Single-cell sequencing data has been deposited at the European Genome-Phenome Archive (EGA), which is hosted by the EBI and the CRG, under accession number EGAS00001004661.

## Acknowledgements

This work received financial support from NCT 3.0 Extension Programm (NCT3.0_2015.17 PrecO-Panc), DKFZ-HIPO (# HIPO-015 K20), PANC-STRAT - single cell sequencing extension (BMBF 01ZX1605C), the incubator grant sparse2big (# ZT-I-0007), and the German Network for Bioinformatics Infrastructure (de.NBI) (# 031A537A, # 031A537C). S.LB. and O.S. were supported by the National Center of Tumor Diseases (NCT) Heidelberg (NCT3.0_2015.17 PrecO-Panc). Human specimens and clinical data were obtained from the EPZ-Pancobank (Biobank of the European Pancreas Center at the Department of General Surgery, University Hospital Heidelberg) in accordance with the regulations of the tissue biobanks and an approval by the Ethics Committee of Heidelberg University (ethic votes 301/2001, 159/2002, S-206/2011, S-708/2019). Activities of the EPZ-Pancobank were supported by the Heidelberger Stiftung Chirurgie and in part by the German Ministry of Science and Education (BMBF) grants 01ZX1305C, 01ZX1605C, 01KT1506. The EPZ-Pancobank is chaired by Prof. Dr. M. W. Büchler, assisted by Dr. N. A. Giese, Dr. M. Schenk, and M. Fischer, and is a member of the BioMaterial Bank Heidelberg/BMBH (coordinator: Prof. P. Schirmacher, BMBF grant #01EY1701) belonging to the German Biobank Alliance. The authors thank K. Schneider, E. Lederer and K. Ruf for the excellent technical support. The authors thank Martin Sprick for helpful discussion.

## Author contributions

OS, CC and RE conceived the project, acquired funding and provided supervision. SL and JJ planned and performed experiments with support from KJ. TGK performed data analysis and wrote the paper. FWT, OD, NI and AG contributed to data analysis. CSL contributed to survival data acquisition and analysis. All authors discussed the results and reviewed the final manuscript.

---

**Supplementary Figure 1:**
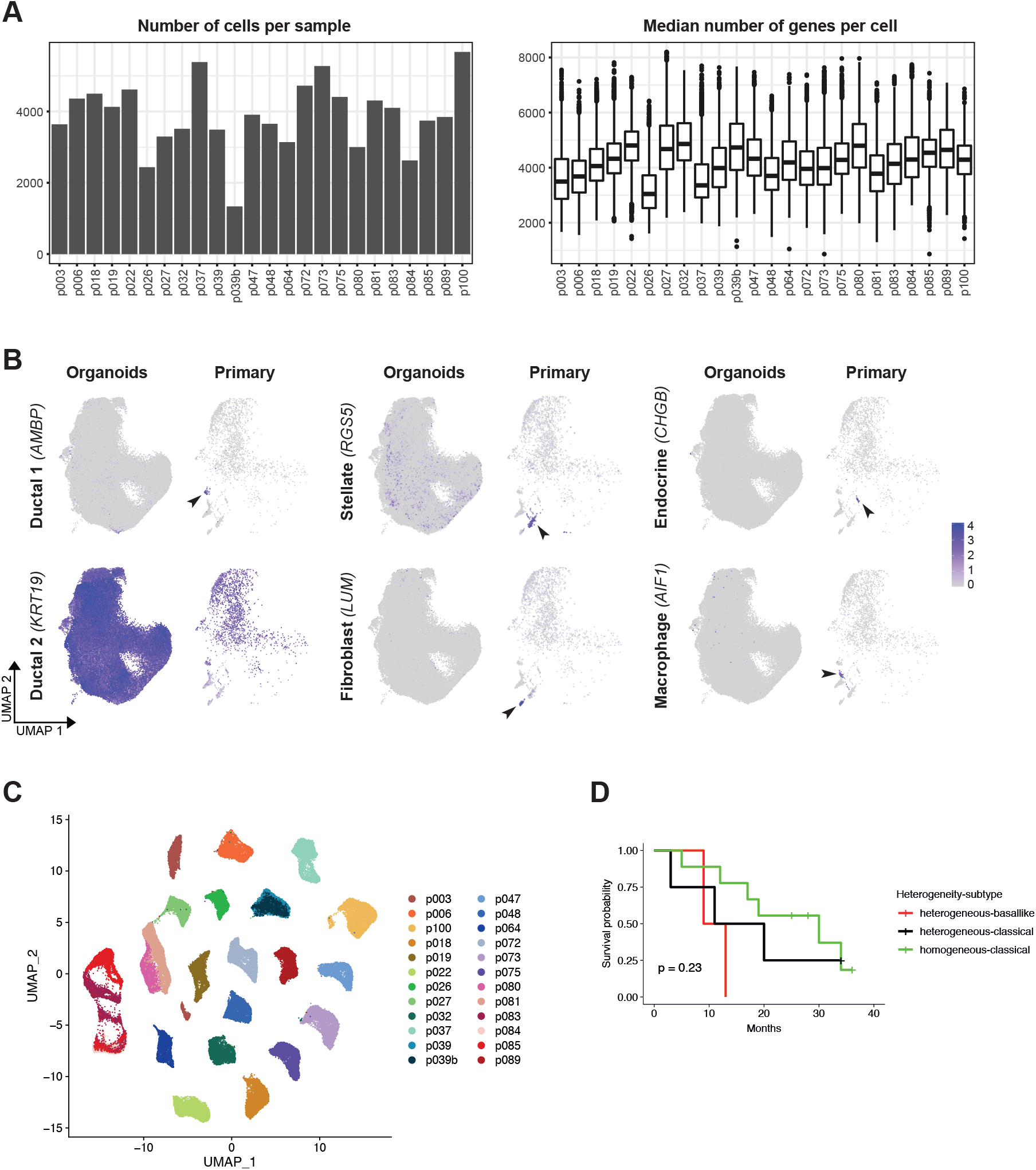
**A)** Number of cells per sample and median number of genes per cell across the PDAC organoid scRNA-seq dataset. **B)** Single-cell expression of cell type marker genes in PDAC organoid cells (left) and cells from primary PDAC biopsies (right). Arrowheads indicate gene expression in small subsets of cells in primary PDAC samples, but not in organoids. **C)** UMAP embedding of 24 scRNA-seq samples from 18 patients, after removal of known expression quantitative trait loci from histologically normal pancreatic tissue samples or PDAC tumor-derived tissue samples ^24^ from the analysis. As also shown in Figure 1C, cells cluster by patient origin (p039 is a technical replicate of p039b, p080 is a biological replicate of p081, p084 and p085 are liver metastases matched to p083). **D)** Kaplan-Meier curve for our cohort, split by PDAC subtype and heterogeneity. Homogeneous classical tumor patients tended to have a better outcome whereas the outcome of basal-like tumors tended to be poorest. While this result is not statistically significant (p=0.23, log-rank test), reflecting that the size of our cohort is insufficient for population-based analyses, it is consistent with previous reports from larger cohorts ^14, 18, 55, 56^.

**Supplementary Figure 2:**
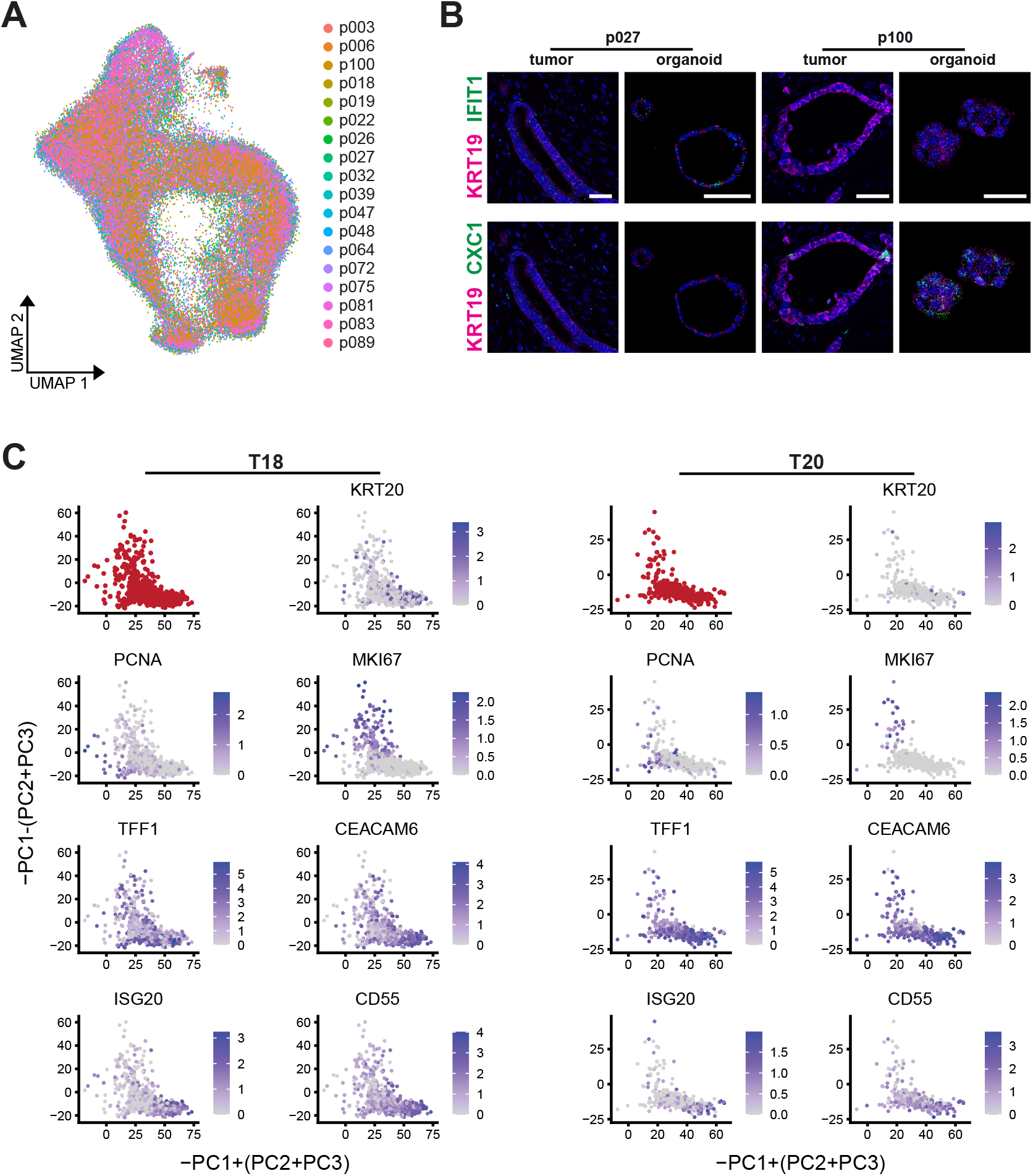
**A)** UMAP representation of PDAC transcriptomes from 18 primary PDAC organoid lines, after PCA-based integration. Colors indicate sample identity. **B)** Representative images of RNA-FISH staining for KRT19 (magenta) together with either IFIT1 or CXC1 (green) in sections from organoids and primary tumours from p027 and p100. Nuclei are stained with DAPI (blue). Scale bars, 50 µm. **C)** Projection of scRNA-seq data from primary PDAC samples T18 and T20, published previously ^19^, onto the same linear combination of principal components as presented in Figure 4F (red). Expression of genes characteristic for cycling and differentiating cells is also shown.

**Supplementary Figure 3:**
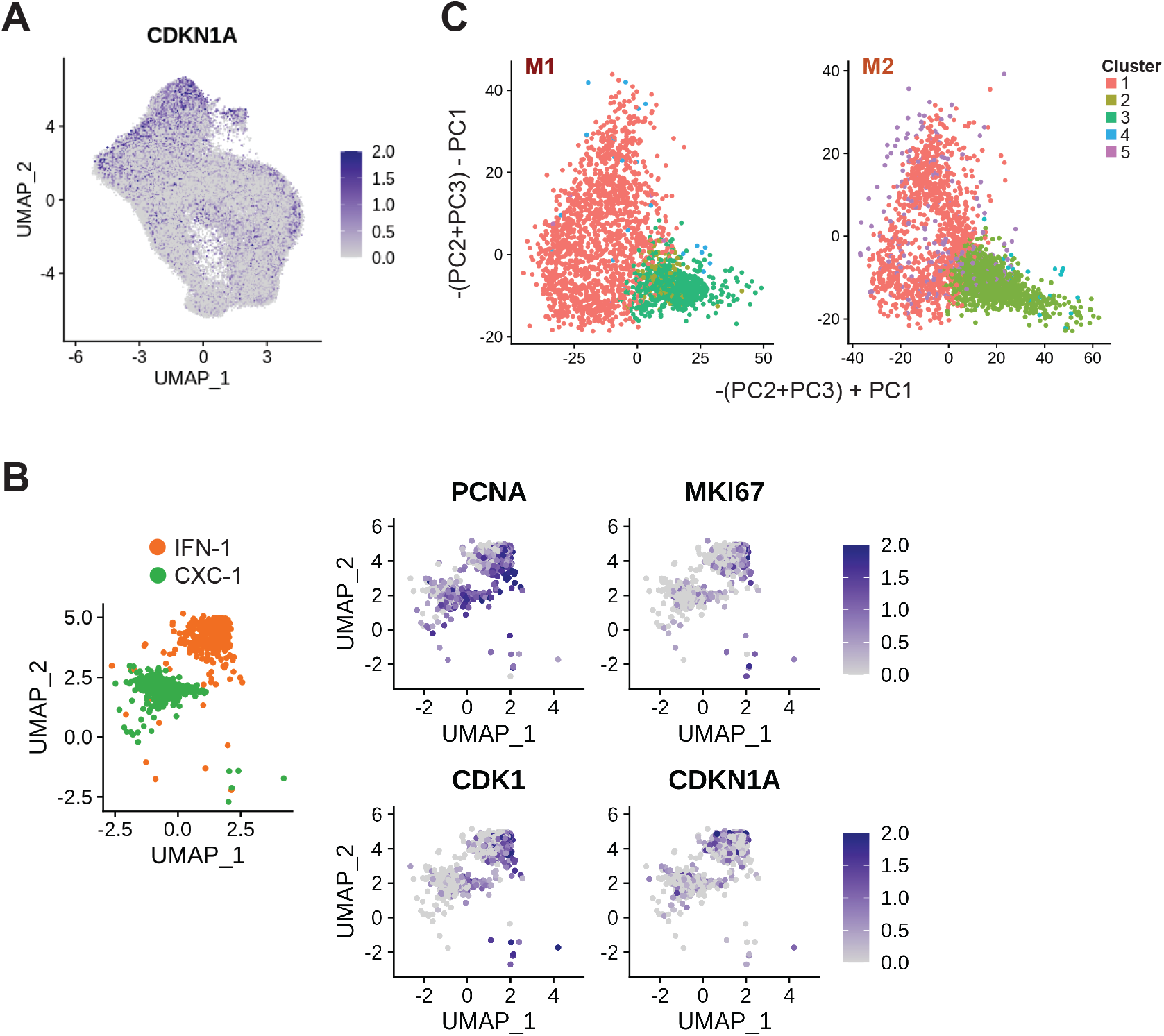
**A)** Expression of CDKN1A (coding for p21) across all primary PDAC organoids. The UMAP representation shown here is the same as in Figure 3C. **B)** For clarity, clusters IFN-1 and CXC-1 are depicted without the other clusters, using the same UMAP representation as above (left). Expression of cell cycle related genes across these clusters indicates that they contain both cycling and quiescent cells (right). **C)** Projection of M1 and M2 transcriptomes onto the primary PDAC organoid data (see Figure 4F and Methods) shows distribution of cells across the functional clusters identified in metastases (see Figure 5A).

**Supplementary Figure 4:**
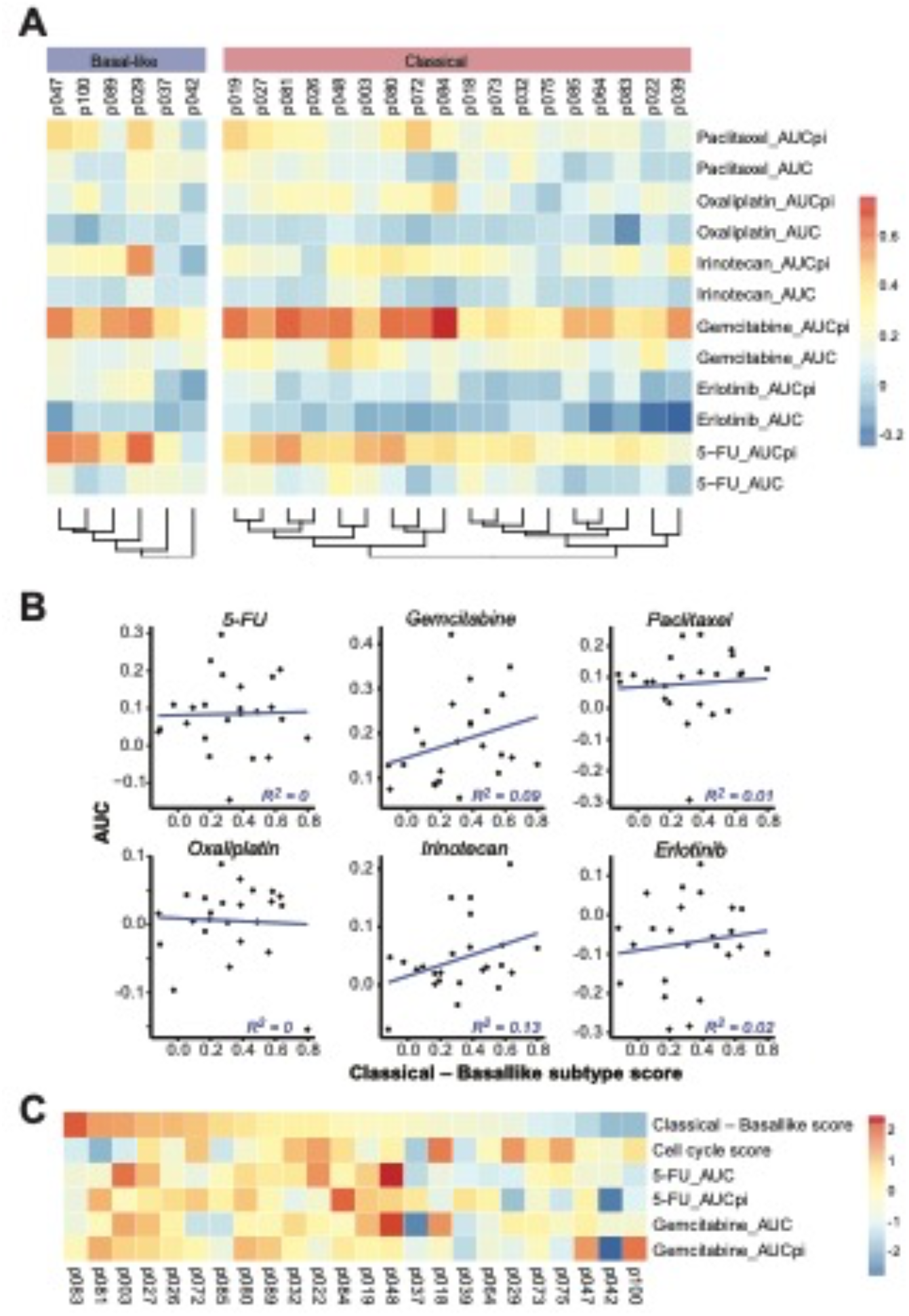
**A)** Drug response as measured by the area under curve for cell death (AUC) and proliferation inhibition (AUCpi) after 72 hours of treatment, for six different drugs in clinical use. Trees represent hierarchical clustering results for ‘classical’ and ‘basal-like’ PDAC organoid lines separately. **B)** Correlation between median subtype scores calculated from scRNA-seq data and drug-induced cell death, as measured by the area under curve (AUC), for 24 PDAC organoid lines. **C)** Overview of variation in transcriptomic PDAC subtype, median cell cycle score, as well as cell death (AUC) and proliferation inhibition (AUCpi) as measured by the area under curve in response to 5-FU or gemcitabine treatment, shown as z-scores scaled across all samples.

## Supplementary Tables

**Supplementary Table 1:** Sample information

**Supplementary Table 2:** Patient-specific genes

**Supplementary Table 3:** Moffitt subtype and heterogeneity scores

**Supplementary Table 4:** Cluster-specific genes

**Supplementary Table 5:** Number of cells represented in each cluster for each patient

**Supplementary Table 6:** Drug screen results

